# Global diversity of integrating conjugative elements (ICEs) in *Helicobacter pylori* and their influence on genome architecture

**DOI:** 10.1101/2025.08.25.671524

**Authors:** Andrés Julián Gutiérrez-Escobar, Mrinalini Srinivasan, Zilia Y. Muñoz-Ramirez, Filipa F. Vale, Difei Wang, Santiago Sandoval-Motta, John P. Dekker, Kaisa Thorell, M. Constanza Camargo, Yoshio Yamaoka, Wolfgang Fischer

## Abstract

Integrating conjugative elements (ICEs) are mobile genetic elements conferring a wide range of beneficial functions upon their bacterial hosts. Generally, they can be activated from their integrated states to undergo horizontal gene transfer via conjugation. In the case of the human gastric pathogen *Helicobacter pylori*, a paradigm for extensive genetic diversity, highly efficient natural transformation and recombination processes may superimpose canonical transfer of its two ICEs termed ICE*Hptfs3* and ICE*Hptfs4*, and thus shape their composition substantially. Here, as a part of the *Helicobacter pylori* Genome Project *(Hp*GP) initiative, we have analyzed high-quality genome sequences from 1011 clinical strains with respect to their ICE content and variability. We show that both elements are highly prevalent in all *H. pylori* populations, but have a strong tendency for gene erosion. ICE sequence variations reflect the population structure and show a clear signature of increased horizontal transfer. A detailed map of ICE integration sites revealed local preferences, but also how recombination processes result in hybrid elements or genome rearrangements. Population-specific differences in ICE cargo genes might reflect distinct requirements in the biological functions provided by these mobile elements.

## Introduction

Infections with the human gastric pathogen *Helicobacter pylori* continue to range among the most common bacterial infections globally, despite a decreasing prevalence in certain countries ^1^, and they represent a major cause for infection-associated morbidity and mortality, particularly due to gastric cancer development ^2^. There is a wide variation in incidence rates between different countries, and a mixture of environmental factors, genetic predisposition of the host, and bacterial virulence factors are thought contribute to disease outcomes ^3,4^. Independently of its virulence factors, *H. pylori* displays a striking genetic variability, with the consequence that virtually all patient isolates differ from each other genetically ^5^. Accordingly, *H. pylori* isolates cluster into geographical populations and subpopulations that have been used to trace human migrations ^6–8^.

While high mutation rates and recombination efficiencies are major contributors to genetic variation, *H. pylori* genomes are also prone to intrusion by classical determinants of genetic variability, such as plasmids, phages and other mobile genetic elements, such as integrating conjugative elements (ICEs) ^9–11^. Generally, ICEs are large elements that are usually integrated into chromosomes, but capable of horizontal gene transfer via self-encoded conjugation systems upon activation, and they often harbor genes encoding antibiotic or phage resistance, metabolic, or virulence traits ^12–14^. In *H. pylori*, ICE genes have originally been described as parts of “plasticity zones” (PZ), clusters of strain-specific genes in a certain chromosomal region ^15,16^. Subsequently, however, comparative analysis of *H. pylori* genome sequences established that these genes are commonly organized as rather conserved genome islands, which can be integrated in these PZ regions, but also elsewhere in the chromosome ^11,17,18^. Two related, but distinct ICE types, termed ICE*Hptfs3* and ICE*Hptfs4*, have been described, and each of them may contain full sets of genes encoding type IV secretion systems (presumably conjugation systems), along with several other genes that partly show a limited conservation between ICE*Hptfs3* and ICE*Hptfs4*, and are partly ICE type-specific ^11,17,19^ (Fig. 1a, 1b). Horizontal transfer of ICE*Hptfs4* between *H. pylori* strains has been demonstrated experimentally, but the canonical conjugative transfer followed by site-specific integration seems to be superimposed by highly efficient natural transformation and homologous recombination processes ^18,20^.

**Fig. 1.**
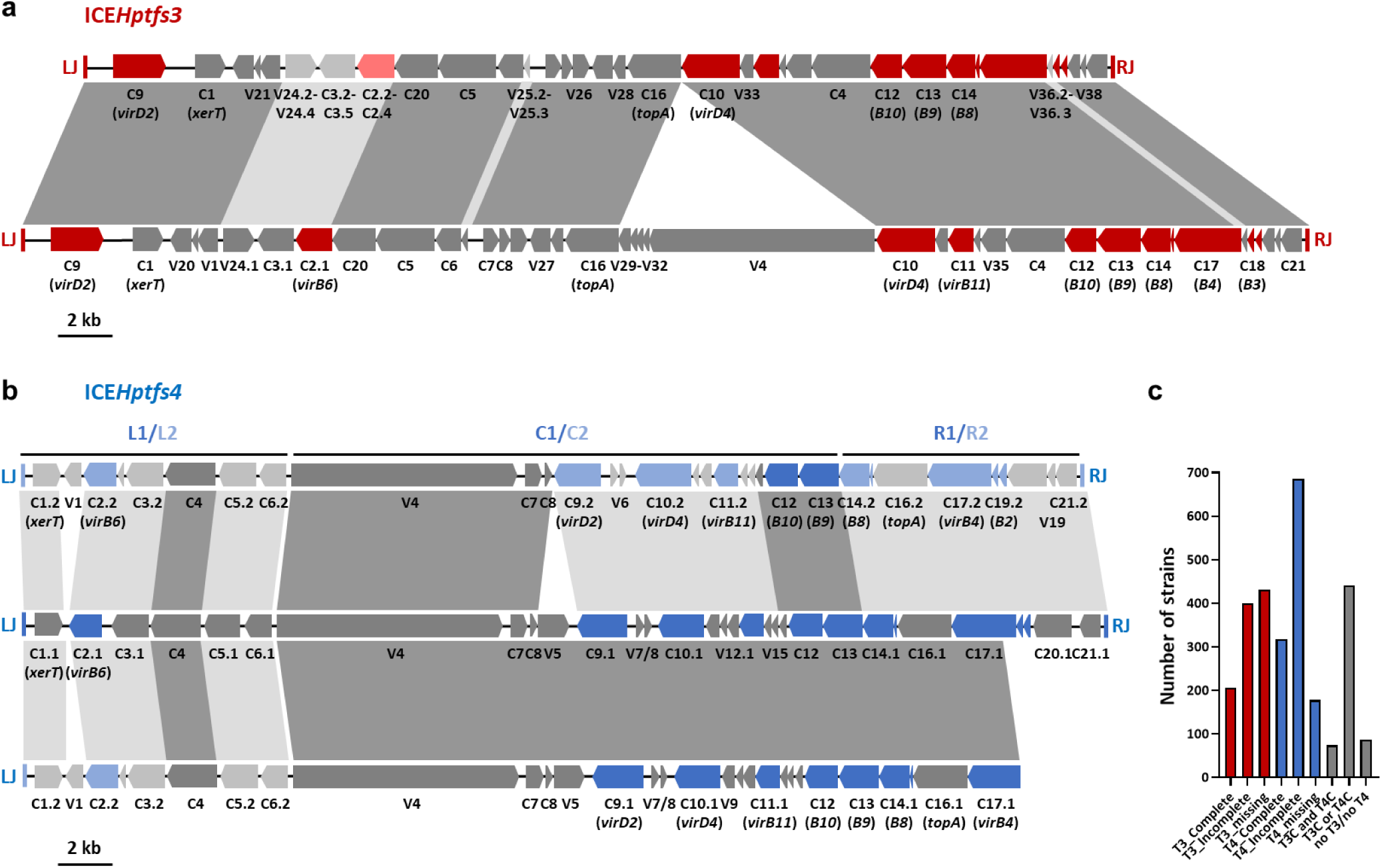
General composition and variants of ICEs in *H. pylori*. (a) Two variants of ICE*Hptfs3* elements are shown, with the respective type IV secretion system (*virB*/*virD*-orthologous) genes colored in red. Variant alleles for individual genes are shown in lighter colors. Light grey sequence similarity bars indicate 25-70% amino acid sequence identity, and dark grey bars 90-99% amino acid identity. (b) Variants of ICE*Hptfs4* elements are depicted, highlighting their modular structure, consisting of distinct versions of left (L1/L2), central (C1/C2), and right (R1/R2) regions. Genes encoding type IV secretion system proteins are colored in blue, and allelic variants are indicated by light or dark shading, as in (a). A frequently occurring, but truncated element, consisting of an L2 left region, a C1 central region, and a truncated R1 right region (here termed R1tr; below), is shown together with full-length L2C2R2 (above) and L1C1R1 (middle) variants. LJ, left ICE junction; RJ, right ICE junction. (c) Numbers of strains containing complete or truncated/incomplete versions of ICE*Hptfs3* (“T3”) or ICE*Hptfs4* (“T4”) elements, or the indicated combinations (“C”, complete; “M”, missing).

Interestingly, correlations between the presence of individual ICE genes (including the duodenal ulcer-promoting gene *dupA*) and gastric disease risk have been indicated in several studies ^21–26^, although some of these correlations turned out to be population-specific ^27^. Furthermore, the presence of certain ICE genes results in higher proinflammatory activity of *H. pylori* ^23,28,29^. For the CtkA (JHP940) protein encoded by one of these genes, a serine/threonine kinase activity towards eukaryotic cells, which affects proinflammatory signaling, has been reported ^30^. Collectively, these previous studies highlight the potential role of ICEs for *H. pylori* pathogenicity.

While the occurrence and properties of ICEs in *H. pylori* strains have been described in a limited number of strains ^11,19^, their full variability has not been addressed systematically. Here, we took advantage of more than 1000 complete, high-quality genome sequences obtained from all over the world in the framework of the *Helicobacter pylori* Genome Project (*Hp*GP) ^31^, to analyze the distribution and properties of ICEs in *H. pylori* isolates on a global scale. The high quality of the genome sequences allowed us to assess not only population-specific features and variations of these elements, but also a systematic analysis of their integration landscape and their influence on genome structure.

## Results

### ICEs are highly prevalent in *H. pylori* genomes, but are subject to truncation or fragmentation

We searched for ICE*Hptfs3* and ICE*Hptfs4* elements within the complete genome sequences of all *Hp*GP strains, using two different approaches: First, left and right ICE junctional regions were identified by BLAST searches, using 80 bp or 200 bp query sequences adjacent to the left and right junctions (see Supplementary Table 1 and Methods for details). In this way, 2609 left or right ICE junctions were identified in the 1011 genome sequences of the *Hp*GP data set. Second, we performed Megablast searches of all *Hp*GP sequences, using three full-length ICE sequences that represent all ICE variants described so far. Overall, we identified 608 ICE*Hptfs3*, and 1002 ICE*Hptfs4* elements, including truncated versions (Supplementary Table 2). Despite a range of variability in the sequences, all regions could be assigned to either ICE type, and no additional or distinct ICE versions containing typical genes with lower sequence similarities were detected.

ICE*Hptfs3* elements comprising a full set of genes had an average size of 46.5 kb, while a substantial percentage of elements specifically lacked the V4 ortholog and a few short open reading frames downstream of V4, resulting in an average size of 38 kb (Fig. 1a). Thus, 160 out of 637 elements contained a full set of ICE*Hptfs3* genes, while the V4 gene, and partly genes V29-V32, were missing in 47 additional elements. Since the latter variant is thus not uncommon, and since ICE*Hptfs4* elements often carry V4 genes with very similar sequences (see below), we considered both variants as complete ICE*Hptfs3* elements (independently of frameshifts in individual genes). The remaining 430 ICE*Hptfs3* elements were truncated to varying degrees, ranging from one or a few missing genes to the absence of major ICE parts including one or both junctional regions (Supplementary Table 2). On average, these truncated, and thus “incomplete”, ICE*Hptfs3* elements had a size of 20.8 kb.

In the case of ICE*Hptfs4* elements, two alleles can be distinguished for most individual genes, and sets of alleles are commonly linked to form corresponding left (L1/L2), central (C1/C2), or right regions (R1/R2), which can be variably combined (Fig. 1b) ^19^. Furthermore, we have previously described a characteristic truncation close to the right junction of R1 regions, in which genes C18-C21, as well as the 5’ end of C17, are missing ^11^ (Fig. 1b). Since the missing genes partly encode essential functions in the corresponding type IV secretion system, this truncated variant cannot be considered (functionally) complete; nevertheless, due to its high prevalence, we considered this version as “complete/truncated” (C/tr) in those cases where all other genes were present. In this sense, 320 out of 1002 ICE*Hptfs4* elements were complete (269) or complete/truncated (51), while the remaining 682 elements lacked at least one, but frequently several or most genes and were termed incomplete (with an average size of 13.7 kb). Among the complete ICE*Hptfs4* elements, almost all possible combinations of left, central, and right modules were found, albeit at different frequencies (Supplementary Table 3). Thus, L1 regions were most commonly found in L1C1R1 combinations (previously also called ICE*Hptfs4b* ^11^), whereas L2 regions were preferentially found as L2C2R2 (previously ICE*Hptfs4c*), but also frequently as L2C1R2 (previously ICE*Hptfs4a*) or L2C1R1 combinations. The R1tr variant was most often associated with L2 and C1, and it seems likely that the L2C1R1tr combination is a fixed combination in certain populations (see below).

When we compared all 1011 *Hp*GP strains, ICE*Hptfs4* elements were more common than ICE*Hptfs3* elements (Fig. 1c), with 311 strains containing complete ICE*Hptfs4* (including C/tr) elements versus 204 strains containing complete ICE*Hptfs3* elements, and 406 strains lacking ICE*Hptfs3*, but only 170 strains lacking ICE*Hptfs4* sequences. A total number of 441 strains (43.6%) have at least one complete element (ICE*Hptfs3* or ICE*Hptfs4*), whereas 483 strains (47.8%) contain fragmented ICEs only. The number of strains completely devoid of ICE*Hptfs3* or ICE*Hptfs4* sequences was thus rather low (87 strains; 8.6%). We found 73 strains with at least one complete ICE*Hptfs3* and one complete ICE*Hptfs4* element, three strains with two complete ICE*Hptfs3*, and six strains with two complete ICE*Hptfs4* elements (Supplementary Table 2). Among the latter nine strains, four strains contained even three complete ICEs, which indicates that there is no active restriction of their number in individual strains (such as an entry exclusion process).

### Strains from all *H. pylori* populations harbor ICE*Hptfs3* and ICE*Hpfts4* elements, but with population-specific preferences

Given the average size of 46.5 kb for full-length ICE*Hptfs3*, and 39.5 to 41 kb for the different full-length ICE*Hptfs4* variants, carrying more than one ICE implies a substantial contribution to genome size. However, due to the frequent truncation of both element types, we found a wide distribution of ICE content in individual strains, ranging from zero to more than 135 kb, or 7.9 % of the full genome size (Fig. 2a). When we plotted the chromosome length of individual strains against their total ICE content, a direct correlation was evident for the complete set of strains, with a mean chromosome size of 1634.4 ± 40.9 kb, and an average ICE content of 39.7 kb (ICE*Hptfs3*, 18.0 kb; ICE*Hptfs4*, 21.7 kb). Similar correlations were also seen for individual *H. pylori* populations with comparatively high or comparatively low average genome sizes (Fig. 2a), suggesting that carrying ICEs does not influence gain or loss of additional genetic material, and that population-specific differences in genome size are not due to differences in the average amount of acquired ICE sequences.

**Fig. 2.**
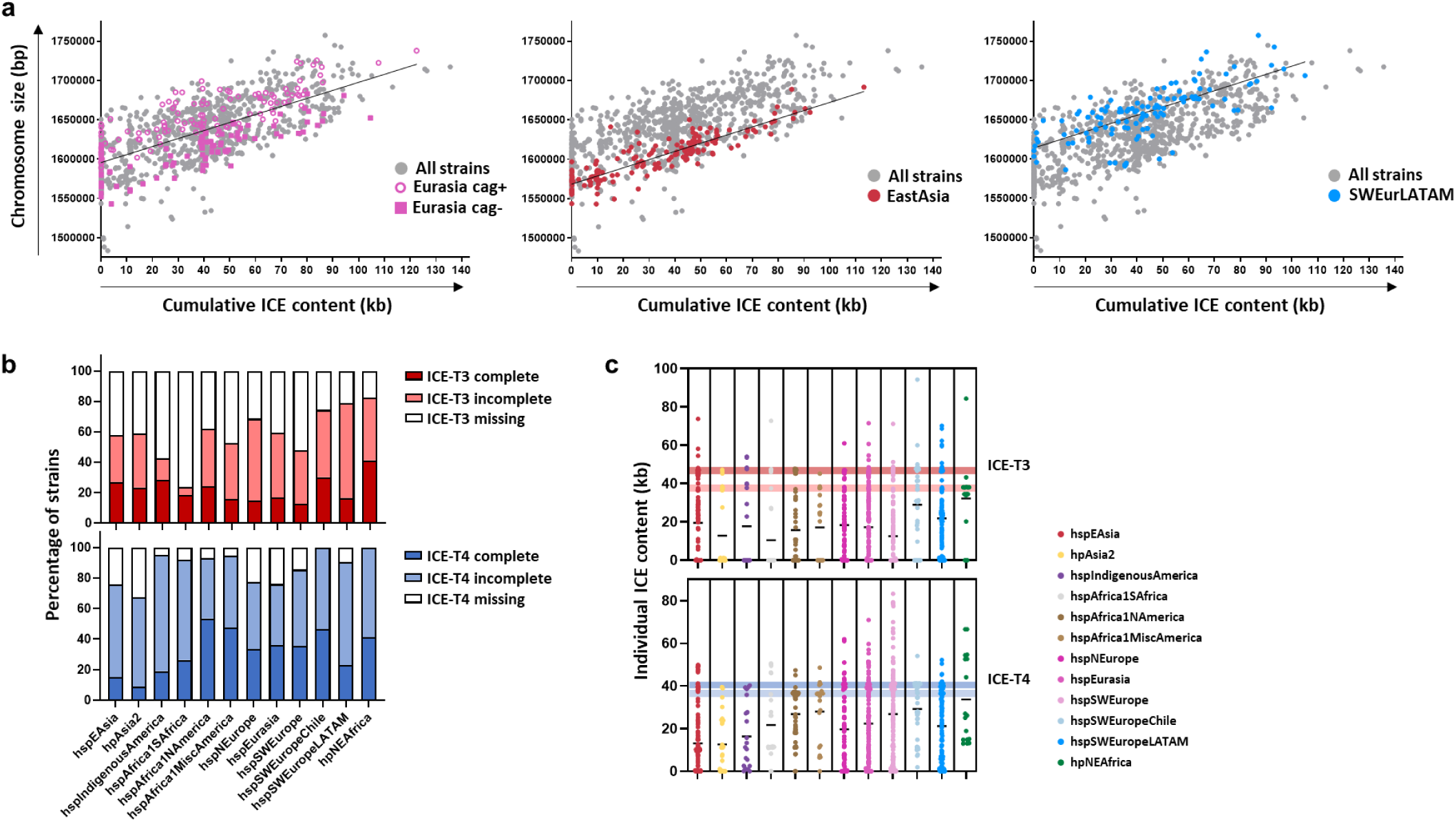
Population-specific distribution of ICEs. (a) Scatter plots of total ICE sequence content (in kb) versus the respective genome sizes for 1011 strains, with highlights for three populations showing either a broad distribution of genome sizes and variable *cag*PAI presence (hspEurasia), smaller average genome size (hspEastAsia), or bigger average genome size (hspSWEuropeLATAM). Note that ICE content does not substantially differ between hspEastAsia (extrapolated average genome size of an ICE-free strain 1,568 kb) and hspSWEuropeLATAM (extrapolated average genome size of an ICE-free strain 1,614 kb) (b) Frequency of strains containing complete, incomplete, or no ICE*Hptfs3* or ICE*Hptfs4* elements in distinct populations. Only populations with at least 16 strains in the *Hp*GP data set are shown. (c) Plots showing the variability of ICE*Hptfs3* or ICE*Hptfs4* content (in kb) between individual strains from different populations. Mean values are indicated for each population. Colored horizontal bars indicate the size range of full-length and shortened ICE versions (ICE*Hptfs3* without V4 and V29-V32, or ICE*Hptfs4* R1tr), respectively.

Although both ICE*Hptfs3* and ICE*Hptfs4* elements were found in all *H. pylori* populations, their relative prevalence differs. Generally, complete or incomplete ICE*Hptfs4*-type elements were more frequent than (complete or incomplete) ICE*Hptfs3* elements in all populations (Supplementary Fig. 1). However, populations hpAsia2, hspEAsia, and hspIndigenousAmerica, which showed the lowest prevalence of complete ICE*Hptfs4* among all populations, had a preference of complete ICE*Hptfs3* elements over complete ICE*Hptfs4* elements (Fig. 2b). These populations also had the lowest average content of ICE*Hptfs4* (Fig. 2c). On the contrary, different hpAfrica1 subpopulations, as well as hpNEAfrica, and the hspIndigenousAmerica and hspSWEuropeChile subpopulations, reach a prevalence of complete or incomplete ICE*Hptfs4* elements close to 100 %, which is also partly reflected in their higher average ICE*Hptfs4* content (Fig. 2b, Fig. 2c, Supplementary Fig. 1). Generally, only very few strains assigned to hpAfrica1, hpAfrica2, or hspSWEurope subpopulations other than hspSWEurope itself, were devoid of both types of elements, whereas a complete absence of ICE*Hptfs3* and ICE*Hptfs4* was more frequent in hpAsia2, hspEAsia and hspEurasia strains. Since the only hpSahul strain in the *Hp*GP data set merely contains a vestigial ICE*Hptfs4* fragment (Supplementary Table 2), a conclusion about ICE prevalence in this population was not possible.

We also checked the ICE content of strains belonging to the *H. pylori* strains that cluster with their respective geographical populations, but form a distinct lineage due to the presence of highly divergent chromosome regions ^32^. All Hardy strains in the *Hp*GP data set belong to the hspIndigenousAmerica subpopulation, and have a low average genome size as a distinguishing feature (Supplementary Fig. 2). Nevertheless, their chromosome sizes vary with their ICE content in the same way as in strains from other populations, even though the maximal ICE content we found among the *Hp*GP Hardy strains was only 42.7 kb. This implies that despite their obviously reduced genome size (with an estimated average of only 1493 kb for an ICE-free Hardy strain; Supplementary Fig. 2), ICEs are still common.

While hpAfrica2 population strain genomes are consistently devoid of the *cag* pathogenicity island (*cag*PAI) ^33^, different other populations are *cag*-positive to varying degrees. For example, the hspEurasia population in *Hp*GP has a frequency of only 56.5 % *cag*-positive strains, as compared to 85.1 % for hspSWEuropeLatinAmerica (hspSWEuropeLATAM) strains, or 96.4 % for hspEastAsia strains. Distinguishing between *cag*-positive and *cag*-negative strains in individual populations, such as hspEurasia or hspIndigenousAmerica (Fig. 2a, Supplementary Fig. 2), or over all strains (Supplementary Fig. 3), showed that their distribution of ICE content is independent of their *cag* status, while the presence of the *cag*PAI is also reflected in higher average chromosome sizes in comparison to *cag*-negative strains. There was no significant association over all populations between ICE carriage and the presence of the *cag*PAI (Chi-square test, p = 0.076). Thus, ICE content and *cag*PAI presence are independent major contributors to genome size in individual strains.

In order to estimate the congruence of ICE sequence variations with core genome variations, we took advantage of the V4 genes, which are sufficiently conserved between ICE*Hptfs3* and ICE*Hptfs4* elements to allow comparative analyses, and used 514 V4 sequences from both types of elements to perform distance network analysis. The results show a network which is roughly similar to the core genome network ^31^, except that strains from the hspSWEurope subpopulations are much more scattered than in the core genome analysis (Fig. 3a). Furthermore, strains from these populations, particularly the hspSWEuropeChile and hspSWEuropeLATAM subpopulations, are more strongly connected to hspIndigenousAmerica and hspEastAsia sequences. Strains from the hpAfrica2 population also seem to be more strongly connected to hspAfrica1SAfrica strains and not as isolated as in the core genome analysis. This may reflect the presumably enhanced horizontal transfer rates of ICEs, as compared to core genome sequences in the corresponding geographic regions. To corroborate and extend these findings, we subsequently performed Chromosome painting analysis with the same V4 sequences and 95 reference sequences. In accordance with the network analysis results, V4 genes from hpAfrica2 strains show much more admixture with hspAfrica1SAfrica sequences than in the core genome, and conversely, many hspAfrica1SAfrica strains show substantial admixture from hpAfrica2 (Fig. 3b).

**Fig. 3.**
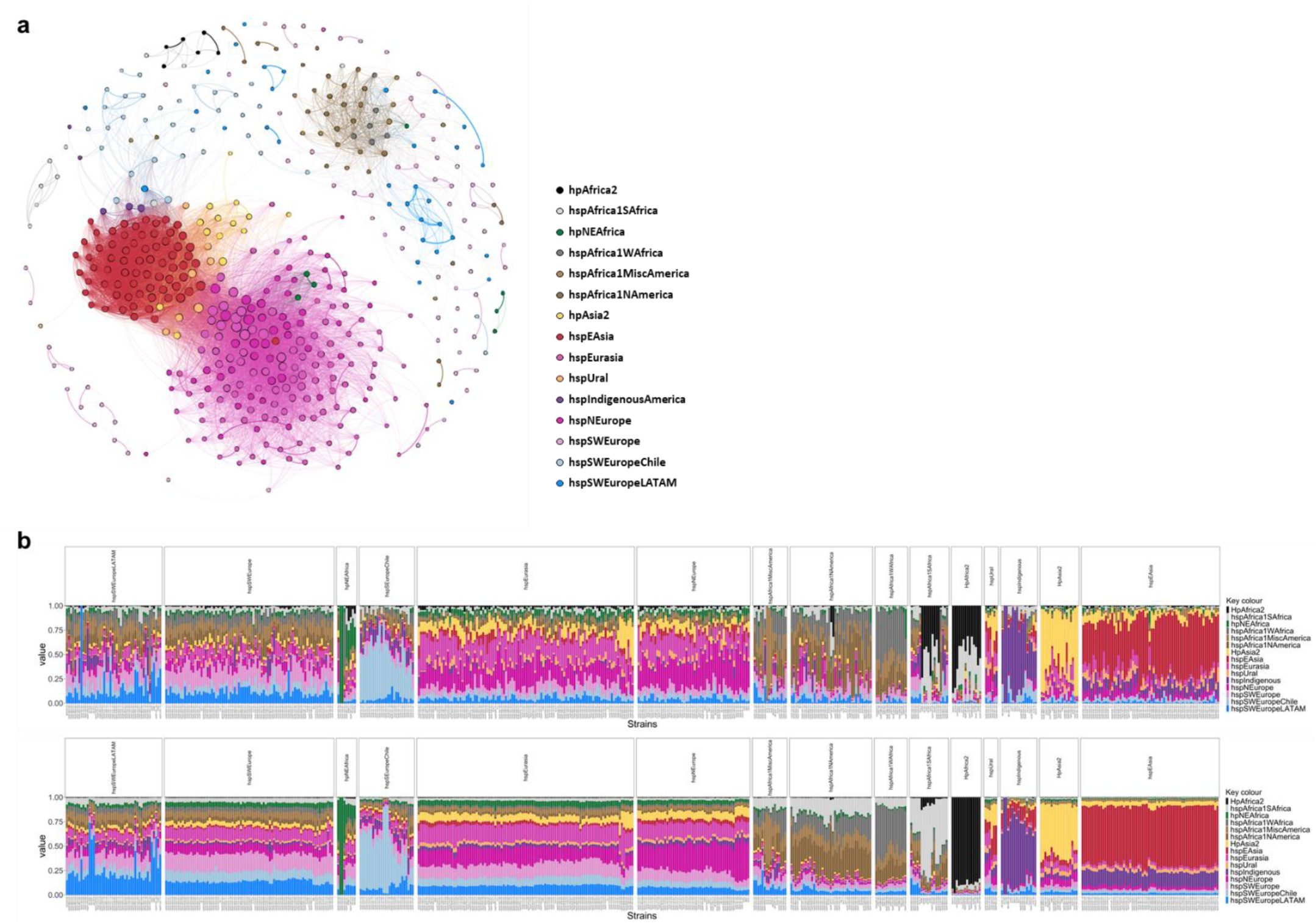
Distance network and Chromosome painting analysis of ICE sequences. (a) A distance network is shown for V4 gene sequences from ICE*Hptfs3* and ICE*Hptfs4* elements. Colors of the nodes represent the populations of the corresponding strains, as indicated. The genetic distance between individual nodes is indicated by the distance and opacity of the connecting lines, and the size of nodes is proportional to their connectivity. (b) Chromosome painting analysis of V4 genes (above) and core genome sequences (below). The vertical bars for each genome represent fractions of ancestral populations and subpopulations in the corresponding sequences, as indicated by the colors.

Several strains from the hpNEAfrica and hspAfrica1NAmerica populations display comparatively strong hpAfrica2 admixtures as well. In fact, strains from most other populations have a stronger hpAfrica2 signature in their V4 genes than in the core genome, supporting the above observation that ICE sequences from hpAfrica2 strains are less isolated than their core genomes. Likewise, many strains from the American hspSWEurope or hspAfrica1 subpopulations, and also strains from other subpopulations that were isolated in the Americas, have much higher hspIndigenousAmerica admixture rates in their V4 genes than in the core genome. Conversely, hspIndigenousAmerica strains often have more admixture from these subpopulations in their V4 genes, whereas admixture from hspEastAsia is more pronounced in their core genome (Fig. 3b).

### Integration sites of ICEs are frequently located in regions of genome plasticity

Comparative analysis of a limited number of complete genome sequences had indicated previously that ICE*Hptfs3* and ICE*Hptfs4* commonly target a short sequence motif (AAGAAT(G)) for integration, and that the presence of such motifs might be sufficient for ICE insertion, without more specific attachment or integration sites required ^11^. To re-analyze this conclusion in the much more comprehensive data set available here, we identified the genes flanking the LJ and RJ sequences for each (complete or incomplete) ICE in which both junctions were present, and mapped the integration sites on the genome of an ICE-free, but *cag*-positive reference strain (HPAG1). In total, 879 ICEs had congruent left and right flanking regions, and could thus be used for mapping (Supplementary Table 4). A further set of 90 ICEs either had congruent left and right flanking regions, but could not be mapped due to the fact that their integration sites are not present in HPAG1 (such as ICEs, PZ regions, or prophages; n=48), or left and right flanking regions were incongruent, i.e. the ICE had integrated into a genome rearrangement site (n=42). Overall, both types of ICEs were found in many different integration sites distributed over the entire chromosome, sparing only a few more extended regions (Fig. 4a). Although we found only one type of element in many distinct insertion sites (84 sites used by ICE*Hptfs3* only, and 99 sites used by ICE*Hptfs4* only), a substantial number of sites (n=50, containing 61.3 % of all insertions) were commonly used by both types (Fig. 4a, Supplementary Table 4). Interestingly, none of the ICEs containing both left and right junctions was integrated in the *cag* pathogenicity island (*cag*PAI) (Fig. 4a, boxed). However, we did find a truncated ICE*Hptfs3* with its left junction inserted in the *cag*PAI (between the *cagQ* and *cagP* genes) in one out of 679 *cag*-positive *Hp*GP strains (VNM-003).

**Fig. 4.**
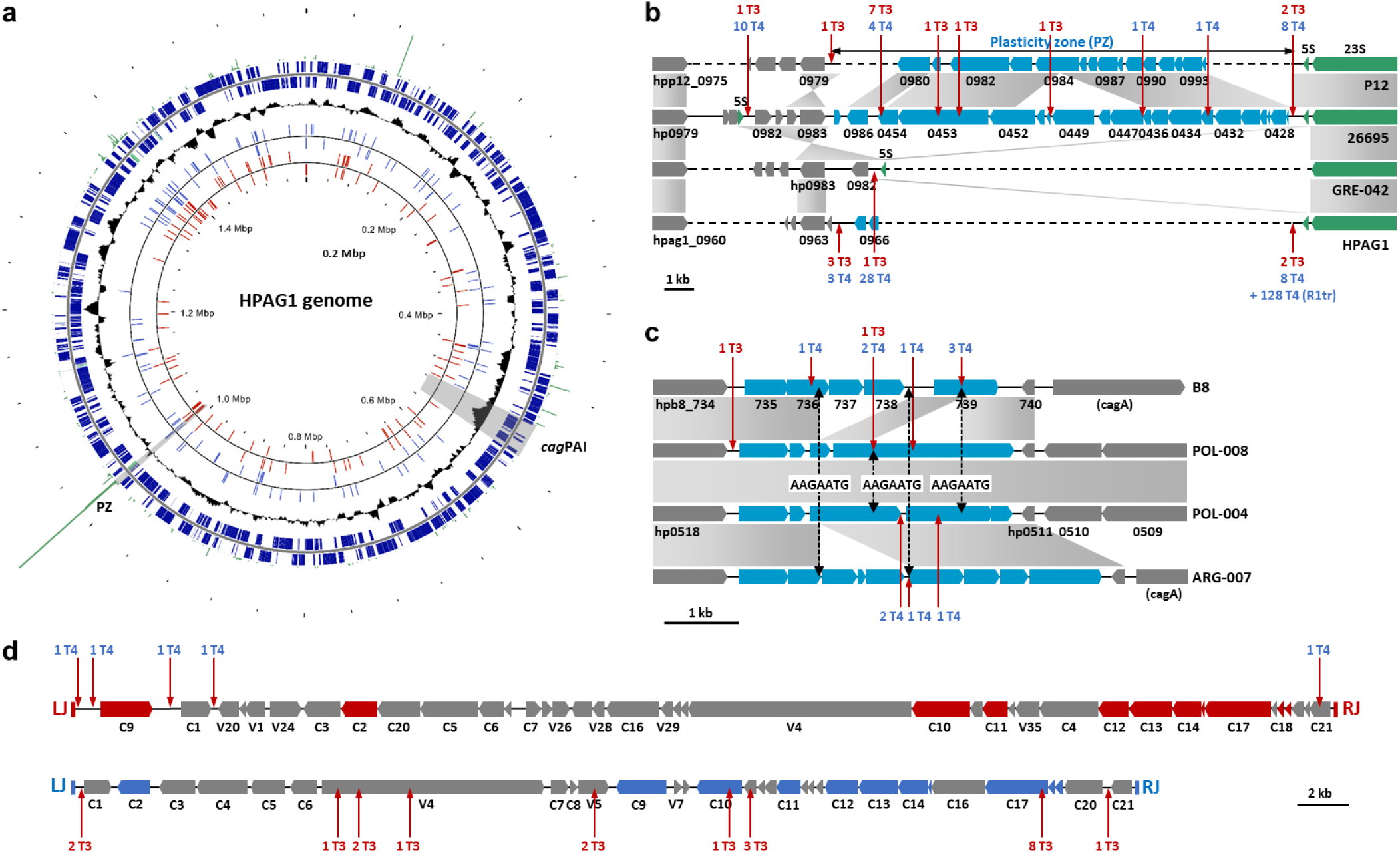
Chromosomal integration sites of ICEs. (a) Circular chromosome map of an ICE-free *H*. *pylori* reference strain (HPAG1), with genes on the plus and minus strands indicated as dark blue bars, and ribosomal RNA genes as green bars (outer circle). The middle circle (black) represents the GC content. Insertion sites of 292 ICE*Hptfs3* (red lines) and 587 ICE*Hptfs4* elements (blue lines) from 1011 *Hp*GP strains (Supplementary Table 4) were mapped onto the HPAG1 genome, and are each shown in clockwise and counterclockwise orientation with respect to their left and right junctions (inner circles). The frequency of insertion at each site (number of ICEs inserted at the corresponding site) is indicated by the green bars outside the map. Localization of the *cag*PAI and the PZ region is indicated by grey shading. (b) Linear maps of representative extended (strains P12 and 26695) and minimal (strains GRE-042 and HPAG1) plasticity zone (PZ) regions. Integration sites and frequencies of ICE*Hptfs3* (T3) and ICE*Hptfs4* (T4) elements in extended and minimal versions of this region are shown by vertical arrows. Note that the PZ region of strain 26695 is shown without its ICE fragments and IS elements (see Supplementary Fig. 4 for further details). (c) Linear maps of a chromosomal region close to the *cag*PAI insertion site, with ICE integration sites and frequencies shown by vertical red arrows, as in (b). AAGAATG insertion motifs at synteny breakpoints are indicated by black dashed arrows. (d) Identified insertion sites of ICEs within previously present ICEs are depicted as in (b). ICE insertions into left or right junction sites, including ILJs, of other ICEs are not shown here.

Some integration sites are used frequently, while others are rare, contributing to a non-random distribution of integration sites (Fig. 4a). The most preferred ICE integration site (211 out of 969 ICE insertions; Supplementary Table 4) is the chromosomal region between the *hp0983* gene and one 5S-23S rDNA locus, which has been termed PZ after genome sequence comparison of strains 26695 and J99 ^15^. In these two strains, the PZ region additionally contains fragments from two or more ICEs ^11^. A closer inspection of PZ regions in *Hp*GP strains indeed revealed a high variability, but also population-specific differences. While strains from the hpAfrica2 population, or from hspEastAsia and the different hpAfrica1 subpopulations, consistently contain a very limited number of genes, or no genes at all, in this region (Supplementary Fig. 4a), hspEurasia, hspSWEurope and hspIndigenous subpopulation strains frequently contain more extended sets of genes there (up to 16 kb of DNA; Supplementary Fig. 4b). Both in genomes with minimal PZ content, and in those with extended PZs, ICEs of either type are most frequently integrated into a site downstream of the corresponding 5S rRNA gene copy, which can be arranged in different orientations with respect to other genes in the PZ, depending on the strain (Fig. 4b). Interestingly, all ICE*Hptfs4* elements containing the R1tr truncation variant (Fig. 1b) are integrated at this position in an otherwise minimal PZ environment (Supplementary Fig. 4a; Supplementary Table 4). In addition to this specific site, we also found examples of ICE integration in several other sites in both the minimal and extended PZ regions, including the sites known for the 26695 reference strain (Fig. 4b, Supplementary Fig. 4b).

Other preferred ICE integration sites include the region close to the second 5S-23S rDNA copy (downstream of the *rbn*/*hp1407*gene), the *hp0088*/*hp0089* intergenic region (which is associated with a genome rearrangement frequently present in hspEastAsia strains), several restriction/modification system genes, another region of genome plasticity (Fig. 4c), which is often found close to the *cagA* gene in strains with a rearranged *cag* pathogenicity island ^34^, but is also often present in *cag*-negative strains ^35^, and other ICEs (Fig. 4d). Similarly to the PZ, the region close to the *cagA* locus contains variable sets of open reading frames that may be heterogeneously fragmented in some, but fused in other strains, including (an) open reading frame(s) encoding an AAA ATPase domain. We found ICE integration within this region in 13 strains, 6 of which are *cag*PAI-positive. Surprisingly, while all ICE insertion sites feature the consensus AAGAATG integration motif, we found that the same motif also delimits certain synteny breakpoints in this region even in the absence of an ICE insertion (such as in strains B8 or POL-008). This suggests that XerT recombinase activity of ICEs (inserted locally or within the same genome) might occasionally shape such (non-essential) regions of genome plasticity by excision of AAGAATG-delimited chromosomal fragments. Re-analysis of the PZ region showed that some synteny breakpoints in this region are also delimited by ICE integration motifs (Supplementary Fig. 4b), collectively indicating that part of the DNA plasticity in these regions has been caused by ICE integration and/or XerT activity during evolution.

### ICEs are major drivers of genome rearrangements

As described above, a substantial number of strains harbors more than one ICE, partly even of the same type, although the latter are frequently incomplete. Among the nine strains with at least two complete ICEs (Supplementary Table 5), three contain two complete ICE*Hptfs3* elements (with or without V4 genes), but we did not find any evidence for recombination between them. The other six strains contain two complete ICE*Hptfs4* elements. Notably, these are always of different subtypes (e.g., L1C1R1 and L2C2R2), but in each strain which contained mixed-type ICE*Hptfs4* elements (i.e., L2C1R1 and L1C2R2 subtypes, respectively), inspection of the genomes indicated that recombination between two ancestral ICEs of L1C1R1 and L2C2R2 subtypes, including inversion of the interjacent genomic region, had occurred (Supplementary Fig. 5, Supplementary Table 5). Two such recombination events had occurred via the V4 genes and led to L2C1R1 and L1C2R2 subtypes, while the third had taken place via the C4 genes and generated mixed-type left regions. Thus, recombination events between multiple integrated ICEs are likely to cause genome rearrangements.

Importantly, recombination between ICE*Hptfs3* and ICE*Hptfs4* elements is also possible via their V4 genes. Although these genes were designated as “variable” genes (i.e., not conserved between ICE*Hptfs3* and ICE*Hptfs4*) ^19^, their actual conservation has not been systematically addressed so far. In fact, at least some V4 gene versions from ICE*Hptfs3* and ICE*Hptfs4* share higher levels of sequence similarity than any of the conserved genes C1 to C21 do ^11^, and similarities are potentially sufficient to enable homologous recombination. A mechanism by which recombination between ICE*Hptfs3* and ICE*Hptfs4* V4 genes leads to the formation of hybrid ICEs containing parts of both elements has also been suggested ^11^(Fig. 5a). As the long-read sequences obtained in the *Hp*GP data set can be unambiguously assigned to either ICE*Hptfs3* or ICE*Hptfs4* V4 genes, this data set enabled us for the first time to analyze similarities between two or more V4 genes present within the same strain. To obtain evidence for recombination between ICE*Hptfs3* V4 genes and ICE*Hptfs4* V4 genes, we aligned 514 full-length V4 sequences (not considering frameshift or nonsense mutations), and generated Neighbor-joining trees. Interestingly, this revealed that the V4 genes do not cluster according to the type of their respective ICEs, but rather according to the strain populations (Fig. 5b). In fact, ICE*Hptfs3* V4 genes and ICE*Hptfs4* V4 genes from the same strain mostly had identical sequences, with the only exception of their extreme 5’ and 3’ end regions (Fig. 5c), suggesting that they readily recombine with each other in the same strain.

**Fig. 5.**
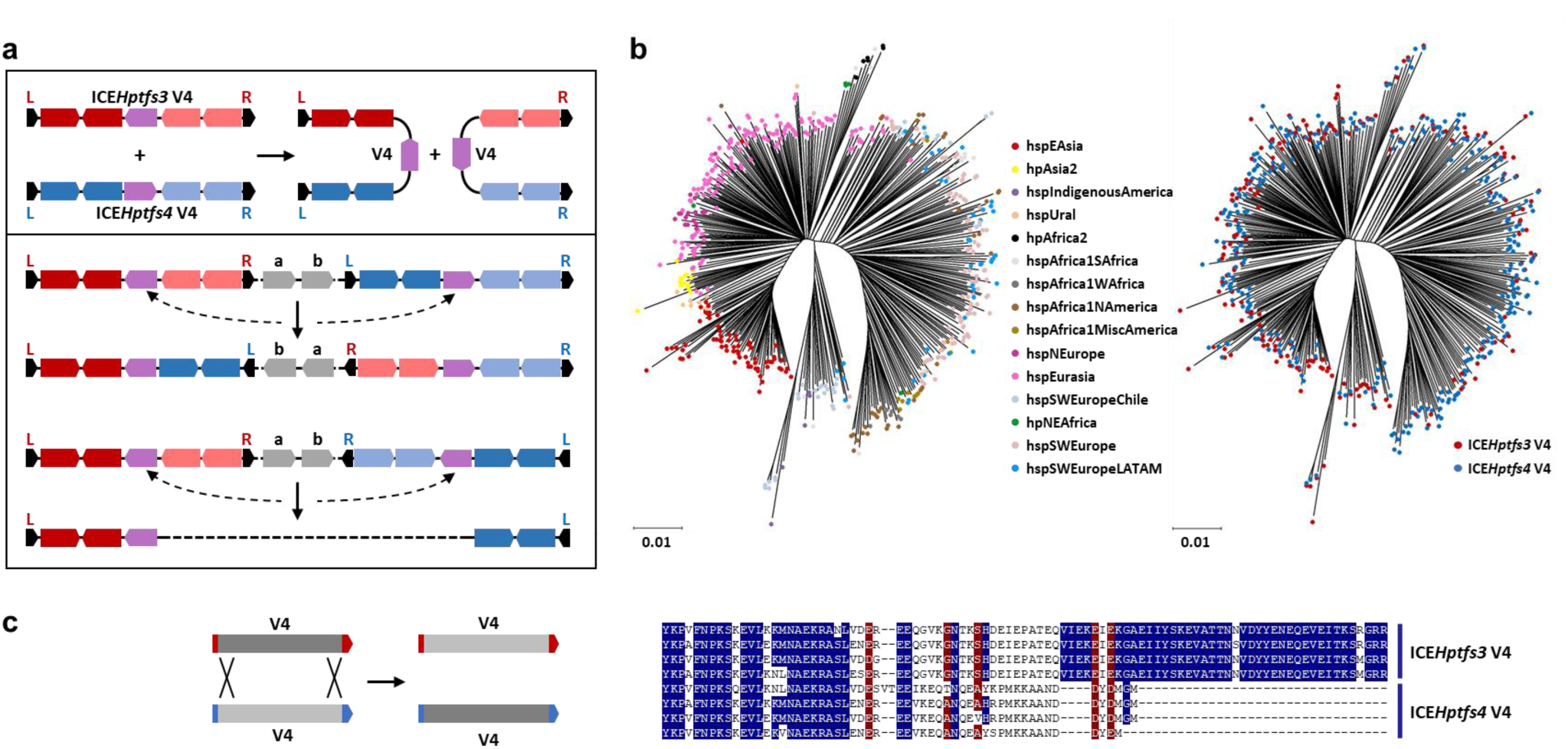
Recombination between ICE*Hptfs3* and ICE*Hptfs4* V4 gene copies and its role for genome rearrangements. (a) Recombination between an ICE*Hptfs3* and an ICE*Hptfs4* V4 gene (purple) generates hybrid ICE versions combining the left or the right regions of each ICE (depicted in stronger and lighter colors, respectively). Such hybrid ICE versions are present in approximately 7 % of all *Hp*GP isolates (Supplementary Table 6). Potential consequences of V4 recombination are concomitant inversions of chromosomal regions, or also deletion of interjacent genes and partial ICEs (lower panel). (b) Neighbor-joining trees of V4 sequences reveal a clustering according to populations (left), but not according to their origin from ICE*Hptfs3* or ICE*Hptfs4* (right). (c) Recombination between ICE*Hptfs3* and ICE*Hptfs4* V4 genes results in exchange and modification of V4 sequences, maintaining distinct N- and C-terminal regions (left). Alignment of representative V4 amino acid sequences shows ICE-dependent C-terminal differences (right).

Next, we identified a total number of 69 strains containing hybrid ICE*Hptfs3*/*Hptfs4* elements in the HpGP data set (Supplementary Table 6). Closer inspection of the resulting gene arrangements showed that 35 hybrid elements resulted from V4 recombination between ICEs inserted independently into the chromosome, and 34 from recombination between ICEs that had inserted into each other (32 ICE*Hptfs3* insertions into ICE*Hptfs4*, 2 ICE*Hptfs4* insertions into ICE*Hptfs3*) (Supplementary Table 6). Most insertions into another ICE had occurred in an antiparallel manner with respect to the left and right junctions (Supplementary Fig. 6a), so that recombination between the V4 copies results in deletion of several ICE genes, including one of the junctions (Supplementary Table 6). The few cases with parallel ICE-into-ICE insertions have usually retained both junctions of the inserting ICE (Supplementary Fig. 6b), although more complex rearrangements (e.g. involving additional IS elements) may have occurred (Supplementary Fig. 6c). Among the chromosomal insertions, 18 V4 recombination events between “parallel” ICEs (Fig. 5a) had resulted in genomic rearrangements (inversions), while no ICE-related inversions could be detected in the other 17 cases (Supplementary Table 6), indicating that double ICE insertion had occurred sufficiently close to each other to prevent deletion of any interjacent essential genes.

### Cargo genes of ICEs have a population-specific prevalence

Both ICE*Hptfs3* and ICE*Hptfs4* elements generally contain conserved sets of genes, with some notable exceptions. As described above, ICE*Hptfs3* elements may lack the V4 gene together with the short open reading frames V29-V32 (Fig. 1a). Furthermore, ICE*Hptfs4* C1 modules additionally contain the V5 gene which is not present in C2 modules, while ICE*Hptfs4* L2 modules additionally contain a V1 gene with substantial sequence similarity to ICE*Hptfs3* V1 genes (Fig. 1b), and to a chromosomal gene that is itself variably present in *H. pylori* strains (e.g., *hpag1_1397*). In addition to these differences, ICE*Hptfs3* elements feature a variable region downstream of the *xerT* recombinase (C1) gene (Fig. 6a). The most common gene arrangement in this region comprises the ICE*Hptfs3* V1 gene, and two overlapping reading frames, one of which (V20) encodes a protein with an AbiEii-like nucleotidyl transferase domain, while the other (V21) might encode an AbiEi-like antitoxin function. Alternatively or in addition to this arrangement, this region may harbor distinct cargo genes, such as the *ctkA* gene, which encodes a protein kinase ^30^, or one of two different genes encoding filamentation induced by cyclic AMP (Fic) domain-containing proteins (Fig. 6a). In the *Hp*GP data set, we identified V20 genes in 482 strains, V21 genes in 497 strains, *ctkA* genes in 113 strains, *fic* (V22.1) genes in 11, and *fic* (V22.2) genes in 53 strains (three of these strains contain both V22.1 and V22.2 on separate ICEs).

**Fig. 6.**
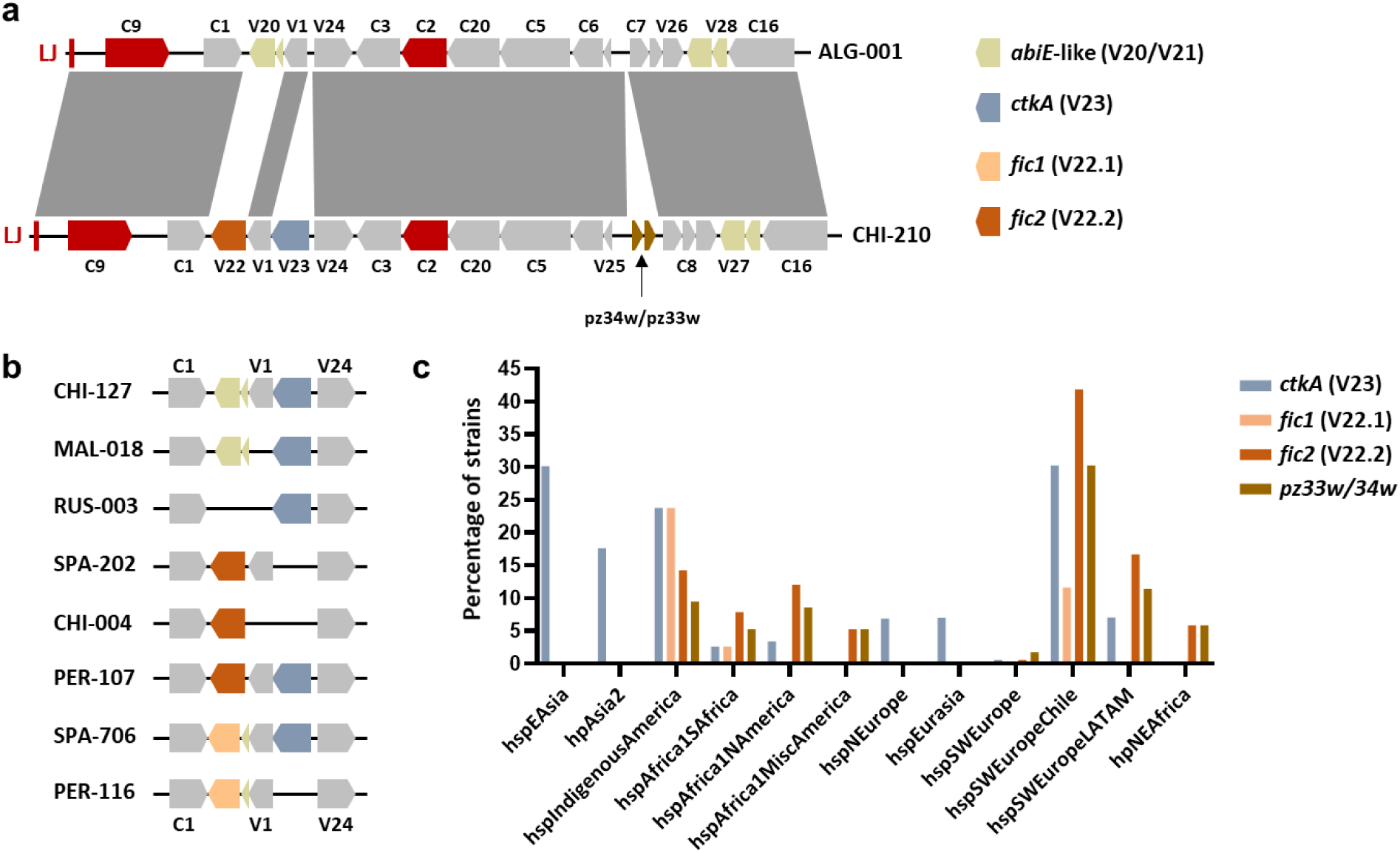
Plasticity and population-specific variants of an ICE*Hptfs3* cargo gene insertion site. (a) Schematic showing the left ICE*Hptfs3* regions of two strains with different cargo gene content. (b) Variability of gene arrangements in the same locus. Note that V22.1 is always in a tandem arrangement with a V21 gene upstream, whereas V22.2 always replaces both V20 and V21. (c) The prevalence of ICE*Hptfs3* cargo genes in *Hp*GP strains from different *H. pylori* populations is shown for cargo genes *ctkA*, *fic1* (V22.1), *fic2* (V22.2), and pz33w/pz34w. Only populations with at least 16 strains in the *Hp*GP data set are shown.

These genes can be arranged with the standard genes in different ways ^29^, but one of the *fic* genes (*fic1*/V22.1) always replaces V20 (i.e., overlaps with the potential antitoxin V21), while the other (*fic2*/ V22.2) is always present without a V21 gene, and thus replaces both V20 and V21 (Fig. 6b). The *ctkA* gene frequently replaces the V1 gene, but can also be inserted additionally between V1 and V24. Two further non-standard genes (or pseudogenes) with similarities to the pz33w and pz34w genes annotated in a 55-kb ICE*Hptfs3* variant of *H. cetorum* (EU015081.1) are inserted between the V25 and C7 genes (Fig. 6a) in 40 strains. In contrast to the other ICE*Hptfs3* genes, some of these genes show a clear non-random distribution in *H. pylori* populations. Thus, the *ctkA* gene is frequently present in strains from hspEastAsia, hpAsia2, hspIndigenousAmerica and hspSWEuropeChile populations, but only rarely present or completely absent in the other populations. In contrast, genes V22.1, V22.2 and pz33w/pz34w are completely absent from hspEurasia, hspNEurope, hpAsia2, and hspEastAsia strains in the *Hp*GP data set, but present at variable rates in other populations, with the highest prevalence in hspIndigenousAmerica, hspSWEuropeChile and hspSWEuropeLATAM populations (Fig. 6c).

Finally, we evaluated the presence of ICE*Hptfs3* and ICE*Hptfs4*, and also the presence of these cargo genes, with respect to disease status for the overall data set, and for the major subpopulations. As shown in Supplementary Table 7, we did not find any significant association between the presence or absence of ICEs (or their completeness) and disease categories, even if individual major populations were assessed. Similar results were observed for the *ctkA* and *fic* (V22.2) genes. However, presence of the *fic* (V22.1) gene, although this is a rare event, was strongly associated with both intestinal metaplasia and gastric cancer (Supplementary Table 7).

## Discussion

The important role of ICEs for adaptation of bacteria to different environments has been documented in many systems ^12,13,36,37^, and their evolutionary interactions with other mobile genetic elements has attracted considerable interest in recent years ^14,38^. In *H. pylori*, where only ICE fragments had initially been identified (in the context of PZs ^15^), comparative analysis of full-length ICEs ^11,16–19^ established that two ICE types, ICE*Hptfs3* and ICE*Hptfs4*, can be distinguished. Although both ICE types can be further grouped into subtypes based on allelic variations, or the absence of particular genes or regions, we did not identify any additional, novel ICE types in this study. We also did not find correlations between ICE types or subtypes and disease development in the *Hp*GP data set. Thus, despite the correlation of some ICE genes with strain virulence, and elucidation of molecular activities of a few ICE-encoded proteins (reviewed in ^27,39^), clear mechanisms of action of ICE*Hptfs3-* or ICE*Hptfs4*-encoded proteins that may provide a potential benefit for *H. pylori* have not been described to date.

We show that ICE*Hptfs3* and ICE*Hptfs4* are widespread in all *H. pylori* populations, but are prone to degradation, confirming and extending earlier observations ^11,19^. Such degraded or incomplete ICEs usually lack one or several ICE core genes, which encode DNA-processing or conjugation functions. Moreover, even in the complete ICEs, these core genes are frequently inactivated by nonsense mutations or insertion of IS elements, collectively indicating that there is no strong selective pressure on maintaining conjugative ICE transfer competence. However, this does not imply that incomplete ICEs are unable to spread horizontally, as they can be efficiently transferred in a non-standard way, independently of conjugation, and involving homologous recombination ^20^. Although we describe certain variations, strains from all populations and subpopulations (except possibly hpSahul, where further data are required) frequently harbor ICEs of either type. This is also true for the Hardy ecospecies, which probably deviated from the standard (Ubiquitous) ecospecies about 200,000 years ago ^32^, and even for the distinct species *H. acinonychis* ^40^ and *H. cetorum* ^41^. Our network analysis and chromosome painting data provide evidence for more frequent mixing of ICE sequences in comparison to the core genome. Thus, even the very distant population hpAfrica2 is represented with higher admixture rates in ICE sequences of most other populations. In isolates from South Africa, where hpAfrica2 strains are prevalent, hpAfrica2 admixtures in the V4 sequences partially reach very high values. Similar observations were made for V4 sequences of North or South American isolates with hpAfrica1 or hspSWEurope ancestry, which contained more hspIndigenousAmerica admixture in comparison to the core genome, and vice versa. These observations are consistent with more frequent horizontal transfer rates of ICE sequences than of core genome sequences. A directional preference of DNA transfer from hspIndigenousAmerica, as concluded from previous observations ^19,42^, was not clearly evident from our data, but we note that hspIndigenousAmerica ancestry is occasionally dominant in V4 sequences from other populations (for example, hspAfrica1SAfrica), whereas no other ancestral V4 sequences are dominant in hspIndigenousAmerica strains. Both the network analysis and chromosome painting results are also consistent with the previous observation of a generally stronger admixture in Latin American subpopulations ^31^.

The high prevalence of ICEs in *H. pylori* populations, and the obvious absence of an exclusion function to prevent multiple ICE intrusion, suggest that beneficial functions and/or maintenance factors encoded in their cargo gene repertoire, are able overcome the fitness cost associated with their presence. One potential fitness cost of ICE carriage is related to the associated increase in genome size. In the *Hp*GP data set, the maximal genome size of an ICE-free strain amounts to 1650.5 kb, whereas the maximal genome size of a strain containing ICEs (which is furthermore *cag*-positive) is 1757.5 kb. However, despite the intriguing observation that different *H. pylori* populations have different average chromosome sizes (Fig. 2a), this does not seem to influence their ICE acquisition capacity, and neither is the average ICE content the reason for such population-specific differences. ICE content is also not reduced in *cag*-positive strains, and there is no general correlation between the ICE content of a strain and the presence or absence of the *cag*PAI. Nevertheless, certain populations with a lower prevalence of the *cag*PAI (e.g., hpNEAfrica, hspSWEuropeChile) tend to contain ICEs more frequently, while some populations with a high *cag*PAI prevalence (e.g., hspEAsia and hpAsia2) have lower rates of ICE presence. In contrast, populations such as hspAfrica1NAmerica and hspAfrica1MiscAmerica are frequent ICE carriers and mostly *cag*PAI-positive. Apart from the *cag*PAI, other accessory genome elements or regions, such as prophages, or the PZ, may contribute substantially to genome size as well. The prevalence of prophages has been shown to differ between populations as well ^43^, but there is an even stronger population-specific bias in the presence of PZ genes, when this term is used exclusively for non-ICE genes, as done here. Thus, strains from several populations, including hpAfrica2, hpAfrica1, and hspEastAsia, generally seem to be devoid of a typical PZ, and carry only a minimal set of genes in this locus, an observation that is retrospectively evident from earlier microarray analysis data ^40^. For example, hpAfrica1 strain J99 ^15^ harbors rearranged ICE*Hptfs3* and ICE*Hptfs4* fragments, including a partial L2C1R1tr ICE*Hptfs4*, in its PZ region, but contains only one short non-ICE gene (*jhp0916*) there. Strains from other populations often have more extended sets of genes in their PZ loci. We show that both the minimal and extended PZ regions are very frequently used as ICE insertion sites, but it should be noted that not all of these sites are necessarily independent. The most striking example is that all ICE*Hptfs4* elements with the R1tr truncation are integrated in the same PZ position, strongly suggesting that the truncation events leading to the loss of C21 to (partial) C17 genes (and a part of the C1 upstream region) resulted in fixation and vertical transmission of this variant, rather than representing independent integration events. However, subsequent recombination with other types of ICE*Hptfs4* has sometimes modified this arrangement to other subtypes (LmC1R1tr, L2C2R1tr, or even L1C1R1tr, after integration of an additional ICE*Hptfs4*). Notably, this variant is not restricted to hpAfrica1 subpopulations, as concluded previously ^11^, but occurs in the hspSWEurope, hspSWEuropeChile, and hspSWEuropeLATAM subpopulations as well, where it is always associated with the “hpAfrica1-specific” *jhp0914* gene ^40^ in a minimal PZ setting. Both the high prevalence of this variant, and its spread to other populations, suggest a beneficial function for the corresponding *H. pylori* hosts.

Apart from this major integration site, both ICE*Hptfs3* and ICE*Hptfs4* elements show a wide distribution of chromosomal integration sites, consistent with their short integration motifs ^11,18^, which thus occur frequently in *H. pylori* chromosomes (approximately 340 sites per Mbp). Nevertheless, most of the identified integration sites are rarely used, and often we identified only a single strain with a given integration site. Not surprisingly, higher ICE insertion frequencies were mainly found in genomic regions containing non-core (non-essential) genes, including already integrated ICEs, restriction/modification system genes, and other variably present regions. Notably, however, these regions do not include the *cag*PAI. We found only one out of 679 *cag*-positive strains with an ICE*Hptfs3* fragment inserted in the *cag*PAI (and associated with a genome rearrangement). In addition, the location of this insertion between the *cagP* and *cagQ* genes is unlikely to render the *cag*PAI non-functional, similar to a *cag*PAI rearrangement caused by IS element insertion between *cagQ* and *cagS*^44^. Thus, although the *cag*PAI does not seem to contain fewer potential integration sites than the rest of the genome (approximately 480 sites per Mbp in strain P12), it seems to be avoided as a target for ICE integration. This is in contrast to prophages, which are more frequently targeting the *cag*PAI ^43^.

One major result of our analysis is that recombination events between multiple ICEs integrated in the same chromosome have probably shaped *H. pylori* genome structure substantially. We show that such events are not only possible between two ICEs of the same type, but also between ICE*Hptfs3* and ICE*Hptfs4* elements via their V4 genes, which share sufficient sequence similarity for recombination. Thus, recombination between two ICE*Hptfs4* elements, which frequently co-occur in the same strain, accounts for most, if not all, mixed-type ICE*Hptfs4* variants (Supplementary Table 3), while L1C1R1 and L2C2R2 are probably the ancestral module combinations ^19^. On the other hand, we detected V4 recombination events, generating hybrid ICE*Hptfs3*-ICE*Hptfs4* elements, in 7 % of all strains. These events often resulted in ICE rearrangements only (after ICE-into-ICE integration or close proximity of insertion sites), but in a substantial number of cases also in genome rearrangements (Supplementary Table 6). Clearly, such rearrangements are only possible if no essential genes are deleted, but they might additionally be restricted by the location of the replication origin and terminus, favoring symmetrical rearrangements with respect to *oriC*, or by other positional biases of certain genes required for optimal expression ^45,46^. Indeed, many large-scale genome rearrangements in *H. pylori* are roughly symmetrical with respect to the origin-terminus axis (Supplementary Fig. 5, ^18^). Genome rearrangements including inversions close to ICE integration sites, for example a major genome rearrangement observed in strain 26695, have been described earlier ^11,15,47,48^. Interestingly, this particular rearrangement in strain 26695 can partly be explained by recombination between the V4 genes of two hypothetical full-length ICE*Hptfs3* and ICE*Hptfs4* elements that had inserted into an ancestral *hp0454-hp0986* fusion gene (Supplementary Fig. 4b) and the 3’ end of *hp0462*, respectively. However, duplication of a repeat region associated with this rearrangement would require an additional event, potentially using a different mechanism as proposed previously ^47^. Strikingly, the intergenic region between *hp0356* and *hp1092* orthologs (corresponding to *hpag1_0352* and *hpag1_0353*), which represents the other breakpoint in this rearrangement, is also a frequent ICE insertion site (Supplementary Table 4). Apart from their potential for genome rearrangements, formation of hybrid ICE*Hptfs3*-ICE*Hptfs4* elements has the remarkable consequence that the canonical junction motifs are no longer available for ICE excision and mobilization (Fig. 5a), due to the respective orientation of the V4 genes. Only site-specific recombination with irregular junction motifs within the hybrid ICE would be feasible in hybrid ICE arrangements, but this is unlikely to result in an ICE variant capable of horizontal transfer. However, it should be noted that even rearranged or truncated ICEs that retain appropriate junctional sequences might still be mobilizable by additional complete elements present in the same bacterial cell, and thus be considered as integrative mobilizable elements ^49^, although such mobilization events remain to be shown.

A further potential consequence of the low integration specificity of the XerT recombinase is that deletions or insertions outside the ICE boundaries may be caused by XerT activity, an assumption supported by the observation of AAGAATG-delimited rearrangement sites (Fig. 4c, Supplementary Fig. 4b). While comparison of integration sites argues that these short motifs are sufficient for XerT site-specific recombination ^11^, it is not clear whether such an activity would be possible without the ICE inserting into the sites to be recombined. Thus, recombinase activity might need additional internal ICE sequences for recognition of the integration site as a substrate. For example, it has been shown that transposition of IS605 family elements requires binding of the TnpA transposase to stemloop structures within the IS element ^50^. Further work is necessary to elucidate the potential impact of ICEs on core genome rearrangements.

Despite their overall conserved gene content, both ICEs are expected to contain cargo genes that are not essential for ICE physiology functions, such as excision, integration, or transfer ^13^. A variable region containing putative cargo genes is particularly evident downstream of the ICE*Hptfs3* C1 gene, where genes from different families have been identified ^11,29^. Several proteins encoded by these cargo genes (CtkA, the Fic proteins and the AbiEii-like protein encoded by V20) share a similar C-terminal region that might contain a secretion signal ^29^. Notably, the ICE*Hptfs3* V27 gene product, which has been shown to encode a further AbiEii-like nucleotidyltransferase ^51^, also contains a similar C-terminal region, while many, but not all, strains have a chromosomal version of the *fic* (V22.2) gene that does not encode this C-terminal extension. The role of these proteins for *H. pylori* fitness, or during infection, is currently unclear, although different studies have provided evidence for an impact of CtkA on human target cells ^29,30,52^. In this regard, the association of the V22.1 gene with intestinal metaplasia and gastric cancer might indicate another candidate for an ICE-encoded host interaction factor, at least in those *H. pylori* populations where this gene is restricted to. However, the number of V22.1-containing strains analyzed in this study is too low yet to clearly draw such a conclusion. Generally, most Fic domain-encoding proteins are AMPylation enzymes, some of which have been shown to be used by pathogenic bacteria as effector proteins for modification of host cells ^53,54^. On the other hand, they might also target bacterial cells, and the genetic organization of V22.1 genes located downstream of, and overlapping with V21 genes supports the notion that they might actually form toxin-antitoxin systems ^54^ involved in ICE maintenance or interbacterial competition, as has been described for the ICE*Hptfs4* V7 and V8 genes ^55^. Future studies are required to determine the biological functions of these cargo genes and to identify their potential role in *H. pylori* pathogenesis.

In conclusion, we provide the most comprehensive and detailed overview of ICEs in *H. pylori* to date. Their long-standing presence in all global populations and co-evolution with their host genomes strongly indicate important functions that remain to be studied, along with their presumable interactions with other mobile genetic elements, such as plasmids or phages.

## Methods

### Data sources

The analysis presented here is based on 1011 genome sequences obtained by PacBio Single-molecule, real-time sequencing in the framework of the *Helicobacter pylori* Genome Project (*Hp*GP), and available at NCBI GenBank under BioProject accession code PRJNA529500. Details of sample acquisition, DNA extraction, library preparation, sequencing and genome assembly have been described elsewhere ^31^. The corresponding genome sequences were used to build an in-house database, and were re-annotated in Prokka v1.14.6 ^56^, using a curated proteome including ICE annotations, as published recently ^19^.

### ICE identification

To identify ICEs within the *Hp*GP genome sequences, we used two different strategies. First, we took advantage of the generally conserved ICE border sequences. Both intact ICE*Hptfs3* and ICE*Hptfs4* are delimited on either side by an AAGAAT(G) integration motif, which is duplicated upon ICE insertion, and followed by conserved sequences depending on the ICE subtype. However, ICE*Hptfs4* elements are occasionally truncated on either or both ends due to irregular recombination at internal AAGAATG motifs ^11^. Therefore, we performed BLASTn searches (using default parameters) with 80-base or 200-base query sequences representing the regular left and right junction ends as well as the typical irregular ICE borders (Supplementary Table 1). Chromosomal positions of ICE insertion sites were listed. Apart from these variants, both types of elements are frequently truncated unspecifically, such that either junction region, or both, are deleted ^11,19^. To identify genome positions of such unspecifically truncated ICE versions independently of the presence of junction regions, and to be able to detect ICEs or ICE fragments of so far unknown types as well, we secondly performed Megablast searches of all *Hp*GP sequences, using three full-length ICE sequences from strain *Hp*GP-ZAF001 that represent all ICE variants described so far, as query sequences.

All identified ICE sequences were subsequently manually curated to determine potential non-ICE elements, such as IS elements or prophages, between the ICE delimitations. Sizes of ICEs were listed without such non-ICE elements, but summing up fragment sizes resulting from disruption due to genome rearrangements. We labeled ICEs as complete (C), if they contained typical arrangements of “conserved” and “variable” genes ^19^. When defined sets of genes were missing, as shown in Fig. 1, ICEs were labeled as “complete, but without a V4 gene” (C w/o V4, for ICE*Hptfs3*), or as “complete/truncated” via irregular right junction (C/tr, for ICE*Hptfs4*). All other ICEs, with deletions ranging from single non-variable genes to most or even all ICE genes, were labeled as incomplete (I). The evaluation of ICEs as complete elements was taken independently of frameshifts of individual genes, or of IS insertion, prophage insertion, or internal gene truncation. The presence of all presumptive (variable) cargo genes (such as V20-V23, V27, V28) was also not considered as required for completeness. For all complete ICE*Hptfs4*, allelic module combinations (subtypes) were also listed. Orthologous groups among complete ICE*Hptfs3* and ICE*Hptfs4* types were calculated using CD-HIT v4.8.1 ^57^. For dot plot representation of individual strains with respect to their genome and ICE sizes, including linear regression analysis, GraphPad Prism v10.2.2 was used.

The presence of the *cag*PAI in each strain was assessed by performing BLASTn with *cagA* 5’ and *cag3* 5’ sequences as query sequences (CP001217.1, positions 585581-586520, and 553525-554528, respectively). Strains were considered *cag*PAI-positive if full-length hits were obtained for both sequences (>87 % identity, and >92.5 % identity, respectively). Association analysis between ICE*Hptfs3* and ICE*Hptfs4* presence and *cag*PAI presence was evaluated using a Chi-square test.

Logistic regression was used to assess the association between ICE*Hptfs3*/ICE*Hptfs4* completeness status and disease categories (172 cases of intestinal metaplasia and 233 of gastric cancer vs. 606 non-atrophic gastritis controls), accounting for variation in ancestry, presence of the *cag*PAI, age and sex. The multivariable models included a revised ancestry variable (8 subpopulations) that collapsed some categories with sparse counts.

### Distance network analysis

Distance network analysis of V4 gene sequences was performed as described previously ^31^. Briefly, 514 V4 sequences, including sequences from two or more V4 copies present within individual strains, were aligned and used to calculate pairwise genetic distances using maximum likelihood criteria in PAUP v4.0a166 ^58^. The resulting distance matrix was normalized to a scale from 0 (highest similarity) to 1 (highest dissimilarity). A fully connected network was then constructed in which nodes represent individual V4 sequences and edges reflect the normalized genetic distances. To resolve the population structure and visualize the relationships between sequences, we applied a pruning approach by sequentially removing the most dissimilar edges until the network separated into distinct connected components (CCs). A CC was defined as a group of at least three connected nodes that are isolated from the rest of the network. Singletons (unconnected nodes) were excluded from the final visualization. The final pruned network consisted of 18 connected components and was colored according to the population assignment of each sequence.

### Chromosome painting

To investigate the ancestry composition of the V4 gene across *H. pylori* populations, we performed chromosome painting. A total of 609 sequences were included: 514 V4 sequences from *Hp*GP strains (including duplicated copies) and 95 reference sequences. Donor strains were defined according to the core genome painting analysis described in ^31^, such that about 15 isolates were selected per population. However, hpNorthAsia and hpSahul populations were not included, since the corresponding *Hp*GP strains lacked V4 genes, and strains with duplicated V4 genes were generally excluded from the donor list (Supplementary Table 8). The analysis was based on a nucleotide alignment of V4 gene sequences, from which SNPs were called using BEAGLE v3.3.2 ^59^. These SNPs were then used as input for ChromoPainter v2 ^60^, which models shared ancestry among recipient and donor populations. Core genome chromosome painting was carried out using the same parameters.

Each recipient sequence was painted using all donor sequences to estimate the proportional ancestry from each population. The results were visualized as stacked bar plots in R, with each color representing the inferred ancestry proportion from a given population.

### Insertion sites

For determination of insertion sites, left and right flanking genes of ICE junctions were noted for all ICEs in which both junction regions were present. When ICE junctions were flanked by other complete or partial ICE sequences, and both ICEs together were flanked by congruent genes, these were noted instead. With the flanking sequences, BLASTn was run against the genome sequences of reference strains HPAG1 (NC_008086.1), 26695 (CP079087.1) and P12 (CP001217.1), and positions of insertion sites were noted for HPAG1, where available. In those cases where both flanking sequences were absent in HPAG1, for example due to insertion of strain-specific restriction/modification system genes in the respective strains, we identified the nearest HPAG1 genes and noted the corresponding contiguous position as an insertion site. For more extended regions that are absent in HPAG1, such as the PZ region, prophages, or ICEs themselves, the flanking regions of ICE insertion sites were mapped directly to these regions. A chromosomal map of strain HPAG1 for visualization of ICE insertion sites was generated using Proksee ^61^.

### Rearrangement analysis and phylogenetic trees

To obtain phylogenetic trees, V4 sequences extracted from 514 ICEs were aligned using the Muscle algorithm within MEGA v5.2 ^62^. Trees were built and tested by neighbor joining with MEGA, using the Kimura 2-parameter model of nucleotide substitution, and 1,000 bootstrap replications. Genome synteny was analyzed and visualized using the Artemis Comparison Tool v7 ^63^ and Easyfig v2.2.5 ^64^.

## Acknowledgements

This work was supported by a grant from the Deutsche Forschungsgemeinschaft (DFG, German Research Foundation) – project number 499103492, to W.F. This research was also supported in part by the Intramural Research Program of the National Institutes of Health (NIH). The contributions of the NIH authors were made as part of their official duties as NIH federal employees, are in compliance with agency policy requirements, and are considered Works of the United States Government. However, the findings and conclusions presented in this paper are those of the authors and do not necessarily reflect the views of the NIH or the U.S. Department of Health and Human Services.

