## Supplementary material for "Global diversity of integrating conjugative elements (ICEs) in *Helicobacter pylori* and their influence on genome architecture": Suppl_Figs_Tables

**Supplementary Figures**

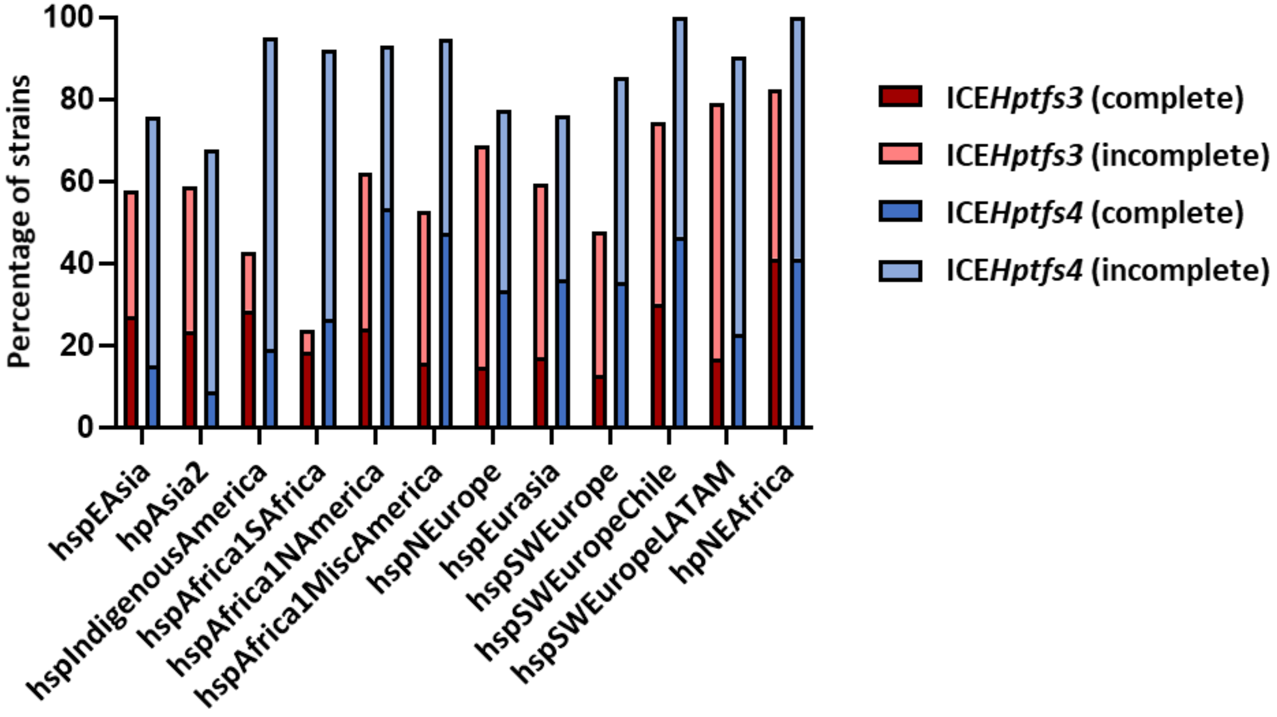


**Supplementary Fig. 1. Prevalence of ICE*Hptfs3* and ICE*Hptfs4* elements in *H. pylori* populations.** Only populations with at least 16 strains in the *Hp*GP data set are shown.

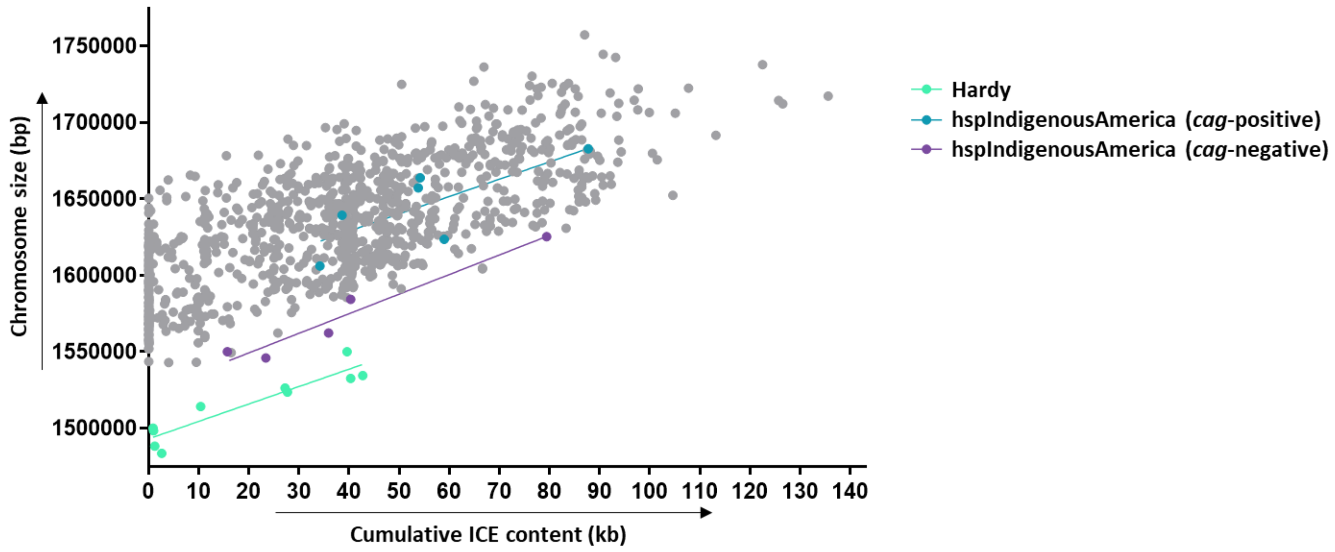


**Supplementary Fig. 2. ICE content in Hardy and Ubiquitous strains from the hspIndigenousAmerica subpopulation.** Ubiquitous hspIndigenousAmerica subgroups are further differentiated according to the presence or absence of the *cag*PAI. Note that all Hardy strains shown here are *cag*PAI-negative.

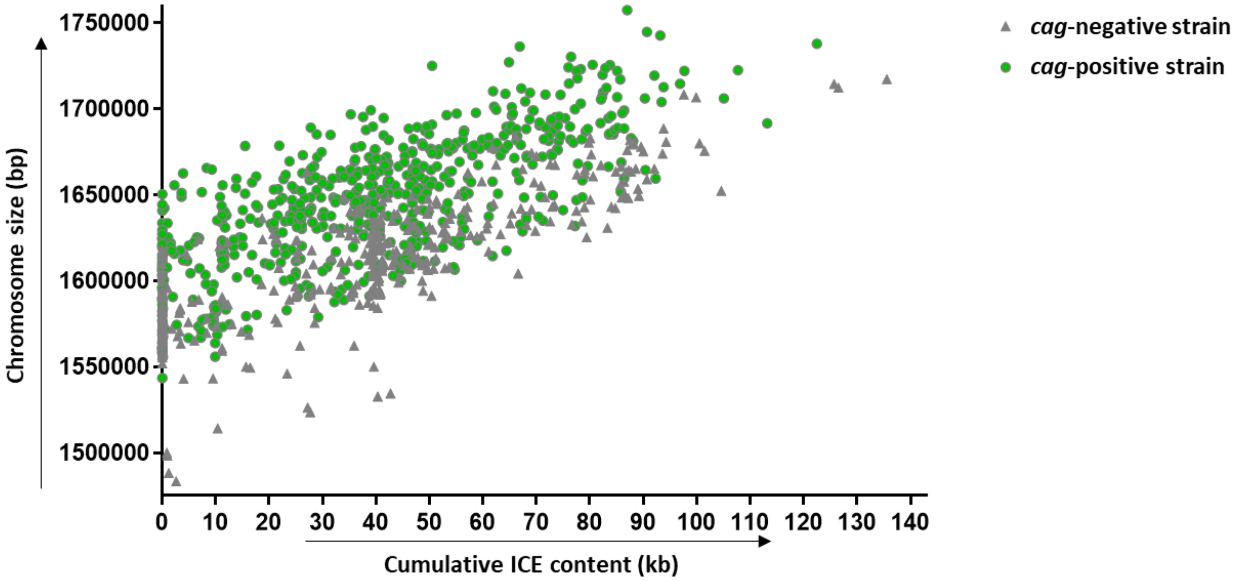


**Supplementary Fig. 3. Distribution of chromosome sizes and ICE content for *cag*PAI-negative and *cag*PAI-positive strains.**

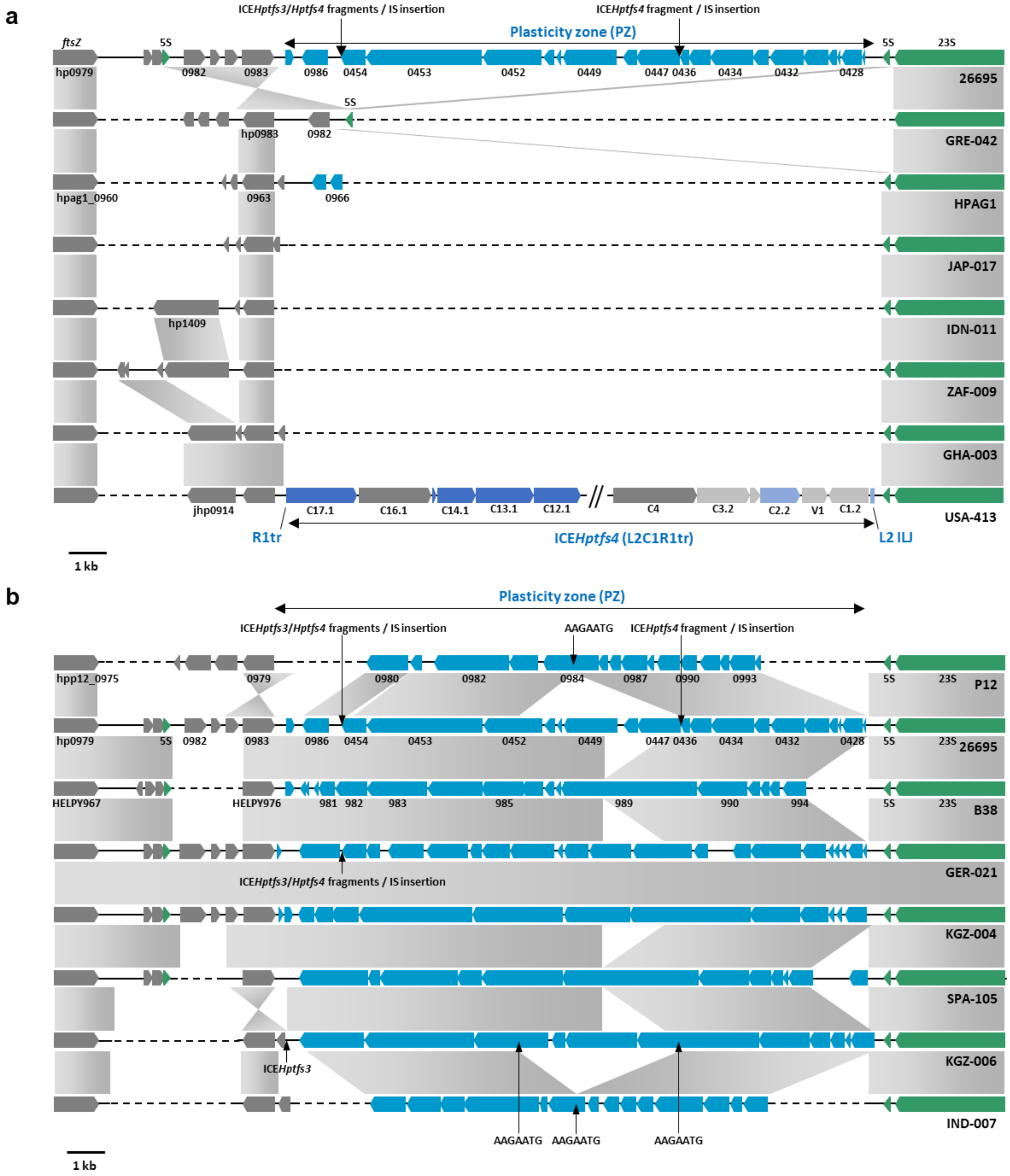


**Supplementary Fig. 4. Variability of plasticity zone regions.** (a) Comparison of plasticity zones (PZ) with minimal gene content with the PZ of reference strain 26695. The PZ gene arrangement of strain 26695 (i.e., genes *hp0428* to *hp0460*, and genes *hp0984* to *hp1009*) is drawn here as one continuous assembly, without the recombination between two copies of a duplicated element (repeat 7) which represents part of the intergenic region between *hp1010* and *hp0462* gene orthologs in the vast majority of *H. pylori* strains. Integration sites of ICE fragments within the 26695 PZ region, including multiple insertion sequences (ORFs *hp0437* to *hp0446*, *hp0455* to *hp0460*, and *hp1009* to *hp0987*) are indicated, but the corresponding ORFs were omitted for clarity. This arrangement thus reflects the consensus PZ gene order in the majority of strains (see panel b below). A subset of strains, including the 26695 reference strain, contain an inversion of a DNA fragment including *hp0983*, resulting in duplication of a 5S rRNA copy. The standard insertion site of ICE*Hptfs4* elements with an R1tr truncation is shown representatively in the minimal PZ region of hspAfrica1NAmerica strain USA-413. (b) Comparison of gene arrangements in PZ regions with extended sets of open reading frames. Integration sites of ICE fragments in 26695 and other strains are indicated, as in (a). Note that open reading frames *hp0454* and *hp0986* in strain 26695 are actually the 5' and 3' regions of one continuous gene in many other strains. Positions of AAGAATG ICE insertion motifs at synteny breakpoints are also indicated. Note that PZ open reading frame delimitations can vary substantially, despite their high sequence similarities.

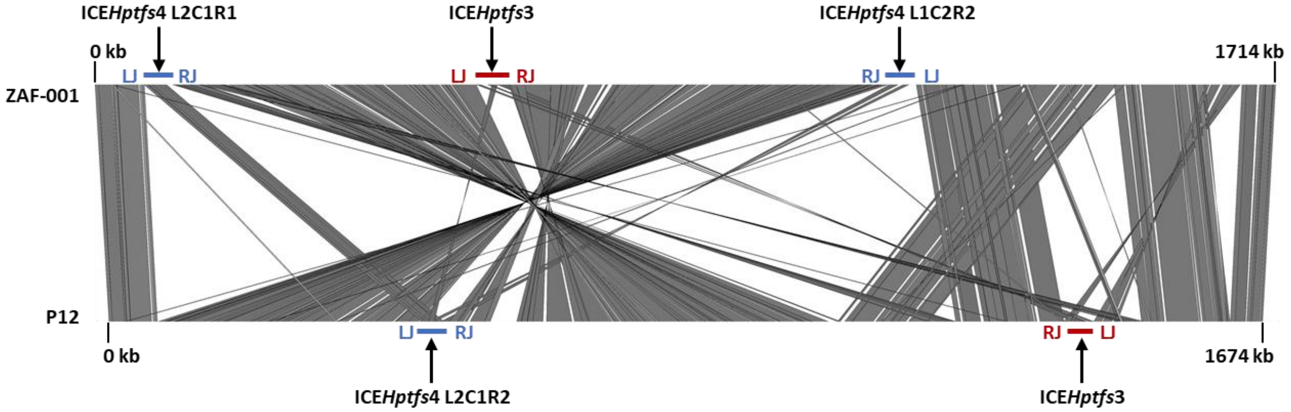


**Supplementary Fig. 5. Chromosomal rearrangement in a strain containing several complete ICEs.** Gene synteny comparison of strain ZAF-001, which contains three complete ICEs, with reference strain P12, showing a large chromosomal inversion presumably caused by recombination between two original ICE*Hptfs4* elements (of subtypes L2C2R2 and L1C1R1, respectively) that were integrated in antiparallel direction.

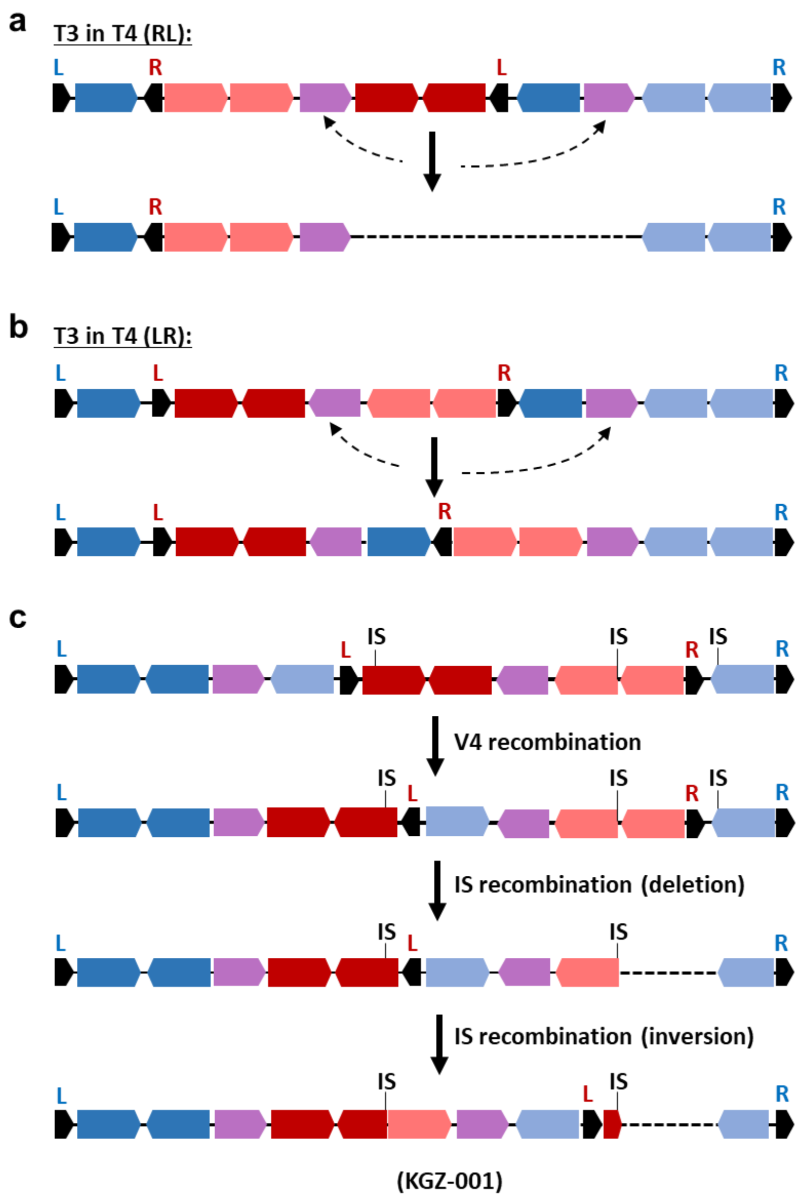


**Supplementary Fig. 6. Possible outcomes of V4 gene recombination between ICE*Hptfs3* and ICE*Hptfs4* elements that have inserted into each other.** (a) Insertion of ICE*Hptfs3* (T3) into ICE*Hptfs4* (T4) in an antiparallel orientation („RL") is depicted. V4 recombination results in deletion of all genes between the two V4 copies, including one ICE*Hptfs3* junction region. (b) V4 recombination between two copies from ICEs that have inserted in a parallel orientation („LR") results in inversion of the interjacent sequence. (c) More complex rearrangements may occur if additional elements, such as insertion sequences (IS), are inserted into ICE sequences. The example shown is from strain *Hp*GP-KGZ-001. Left and right ICE regions, as well as V4 genes, are indicated by colors as in Fig. 5A.

### Supplementary Tables

**Supplementary Table 1. Query sequences used to search for ICE junctional regions.** AAGAAT(G) junction motifs are printed in bold face.

| Designation | Sequence 5'-3' |
|---|---|
| ICE*Hptfs3* left junction (LJ) | **AAGAATG**TAAGTTTTAAGTGTTCTCTAAGTTTCAATCTAAGTCAATTAAAAAATCATTGAAAATGTAGCATAAATTTTTT |
| ICE*Hptfs3* right junction (RJ) | TAGGTTTTAATGTCTTTGCCATAGTTTTCAAGTCCCTTTCTATGAGACTTAGAGACAGAAACTAACATTCTAGT**AAGAAT** |
| ICE*Hptfs4* L1 left junction | **AAGAATG**TAATTATTTAGCTAGCTCTAAGCTTCTTAATTTTCTAATGCTAACTTTTTAAAAACTATTTCTGCGTTCTACC |
| ICE*Hptfs4* R1 right junction | AAGATTTCTACATTAACATCTGGATAGTTTTGGAAGTAAAATTGTTGTGTCTTTTTGGAAAATTTAACATTGTCTAAAAACATTTCTTGGCACTCTTTAGAATTTAAGTGCAGTTTAGGGATTTTAAAAAACTTCATAAACTTCCTTTTAAAAGAAGCTTAGATAAGAGTTCTATAAATAATTACATTCTCTA**AAGAATG**[a] |
| ICE*Hptfs4* L2 left junction | **AAGAATG**GAATTTTTAAGTTAAATCTAATCTACAAAATGATTACTCCGTTGTTATACCAAAAATCTACAAAACTAATCCT |
| ICE*Hptfs4* R2 right junction | AAAACTCTTTCTTGCATGGTCCTAATTCCTAGATATTGATTTCTTAGAGATTAGAAAATAATTGCATTCTCTA**AAGAATG** |
| ICE*Hptfs4* L2 ILJ[b] | **CATTCTT**AACCCTACAAGATGATTTTTATGGATTAAAAATAATTAAGGGAGAAAAATGAGTGATTGCAAAATGAGTAGGG |
| ICE*Hptfs4* L1 ILJ[b] | **CATTCTT**TTTAATTTCTATTAAAAAAATTTAATATTAAGAGAATTTTTATGAAAAAATCAAATGACAATAACGCACTCGC |
| ICE*Hptfs4* R1tr right junction[c] | GTATGGAACTCATCTTATTTAAAATCTCAGCTAAAAGAGTCGCTTTATCTTGCAATTCTTTATTTTCTATTG**CATTCTTT** |

[a] An 80 bp sequence close to RJ is similar to a region adjacent to one 23S-5S-rRNA copy, and would thus be too unspecific.

[b] L1/L2 ILJ, irregular left junction immediately upstream of the respective C1 genes, and in opposite orientation to L1 or L2.

[c] Irregular junction in opposite orientation to R1, and within the C17.1 gene, leading to C17 5' truncation by 183 codons.

**Supplementary Table 3. Distribution of ICE*Hptfs4* subtypes among complete elements.**

| Subtype[a] | Number of elements |
|---|---|
| L1C1R1 | 141 |
| L1C1R2 | 3 |
| L1C2R1 | 0 |
| L1C2R2 | 3 |
| L2C1R1 | 14 |
| L2C1R2 | 30 |
| L2C2R1 | 1 |
| L2C2R2 | 58 |
| L1C1R1tr | 4 |
| L1C2R1tr | 0 |
| L2C1R1tr | 35 |
| L2C2R1tr | 1 |
| LmC1R1 | 11 |
| LmC1R2 | 3 |
| LmC2R1 | 1 |
| LmC2R2 | 3 |
| LmC1R1tr | 11 |

[a] Designation of left (L), central (C) and right (R) regions according to [1], except R1tr, which represents the truncated version of R1 depicted in Fig. 1B. Left regions Lm represent hybrid L1/L2 regions formed via recombination between the respective C4 genes (Fig. 1B).

**Supplementary Table 5. Strains containing multiple complete ICEs.**

| ***Hp*GP strain** | **Type ICE1[a]** | **Type ICE2[a]** | **Type ICE3[a]** | **Recombination between ICEs** | **Inversion from-to** |
|---|---|---|---|---|---|
| CHI-204 | T3 FL | T3 FL | T4 L2C1R2 | none | |
| TWN-027 | T3 w/o V4 | T3 w/o V4 | T4 L1C1R1 | none | |
| USA-423 | T3 FL | T3 w/o V4[b] | | none | |
| COG-002 | T4 L1C1R1 | T4 L2C2R2 | | none | |
| COL-004 | T4 L2(m)C1R1[c] | T4 L1(m)C2R2[c] | | via C4 | ICE internal (C4-C1; C1-C4)[d] |
| ISR-005 | T4 L1C1R1 | T4 L2C2R2 | | none | |
| MEX-008 | T4 L1C1R1 | T4 L2C2R2 | | none | |
| SWT-003 | T3 FL | T4 L2C1R1 | T4 L1C2R2 | via V4 | hp0629-hp0651 |
| ZAF-001 | T3 FL | T4 L2C1R1 | T4 L1C2R2 | via V4 | hpag1_0067-hpag1_1119 |

[a] T3, ICE*Hptfs3*; T4, ICE*Hptfs4*; FL, full length (including V4)
[b] Truncated V4 gene
[c] Left regions are actually mixed (Lm) regions due to the fact that recombination occurred via the corresponding C4 genes.
[d] Both ICEs are inserted in the same site in antiparallel orientation.

**Supplementary Table 7. Association between ICE presence and disease categories (vs. non-atrophic gastritis)**

| | Adjusted odds ratios (95% confidence intervals)[a] | | |
|---|---|---|---|
| | Intestinal Metaplasia | Gastric Cancer | Intestinal Metaplasia and Gastric Cancer |
| ICE*Hptfs3* | | | |
| Missing | 1 | 1 | 1 |
| Complete | 1.06 (0.64 - 1.77) | 0.89 (0.54 - 1.46) | 0.97 (0.64 - 1.46) |
| Incomplete | 0.95 (0.63 - 1.45) | 0.71 (0.47 – 1.07) | 0.82 (0.58 - 1.16) |
| ICE*Hptfs4* | | | |
| Missing | 1 | 1 | 1 |
| Complete | 1.63 (0.89 - 2.98) | 1.39 (0.80 - 2.40) | 1.47 (0.92 - 2.34) |
| Incomplete | 1.58 (0.91 - 2.76) | 1.16 (0.71 - 1.91) | 1.32 (0.86 - 2.03) |
| *ctkA*[b] | | | |
| Absent | 1 | 1 | 1 |
| Present | 0.76 (0.39 - 1.48) | 0.93 (0.50 - 1.76) | 0.85 (0.50 - 1.45) |
| *fic* (V22.1)[b] | | | |
| Absent | 1 | 1 | 1 |
| Present | 12.00 (2.31 - 62.4) | 7.54 (0.81 – 70.0) | 11.28 (2.32 - 54.93) |
| *fic* (V22.2)[b] | | | |
| Absent | 1 | 1 | 1 |
| Present | 2.47 (1.18 - 5.18) | 0.40 (0.11 - 1.45) | 1.47 (0.73 - 2.94) |
| pz33/34w[b] | | | |
| Absent | 1 | 1 | 1 |
| Present | 0.49 (0.17 - 1.43) | 0.24 (0.02 - 2.66) | 0.45 (0.16 -1.29) |

[a] Logistic regression models adjusted for ancestral origin, *cag*PAI, age, and sex.

[b] Restricted to genomes with ICE*Hptfs3*.
